# A comparison of sleep-based and retrieval-mediated memory consolidation using sigma-band activity

**DOI:** 10.1101/2025.08.14.670445

**Authors:** Hayley B. Caldwell, Kurt Lushington, Alex Chatburn

## Abstract

Sleep and retrieval training are reported to promote memory consolidation and are thought to have similar underlying brain-based mechanisms; however, these mechanisms have yet to be directly compared side-by-side, across equivalent timescales. The current study aimed to see if we could replicate sleep spindles’ enhancement of weakly encoded memories, across both sleep-based and retrieval-mediated memory consolidation, using EEG sigma power (∼12–16 Hz) as a proxy for sleep spindles (i.e., sigma-band activity). Thirty subjects (27F, 18–34, *M*=22.17) participated in four separate sessions where they learnt different sets of 104 object-word pairs. Subjects were then tested on their recognition accuracy of the pairs before and after one of four 120-min memory interventions where EEG was recorded: retrieval training (i.e., cued recall practice), restudy (i.e., pair re-exposure), a nap opportunity, or a wakeful rest. Our results did not replicate an enhancement of weakly encoded memories, moderated by either sleep spindles or sigma power. Instead, posterior regions of sigma power negatively related to subsequent memory outcomes. We also detected a prioritisation to enhance memory outcomes for strongly encoded memories and greater memory outcomes following retrieval training and restudy compared to sleep and wake interventions. These results shed new light upon the role of sigma-band activity to predict subsequent memory outcomes and inform how future studies should measure encoding strength and sigma power. Importantly, our study provides a methodological approach to comparing sleep-based and retrieval-mediated memory consolidation that should be explored across different memory paradigms in future studies.

## 1. Introduction

Memory consolidation involves the incorporation of memories into the neocortical landscape for long-term storage (McClelland et al., 1995). An optimal state for memory consolidation is sleep (Rasch & Born, 2013; Staresina, 2024); but recent evidence suggests that memories can undergo rapid consolidation across wake, via repeated retrieval training (i.e., cued recall; Antony et al., 2017; Ferreira et al., 2019; Ritvo et al., 2019). It is assumed that the mechanisms underlying retrieval-mediated consolidation parallel those for sleep-based consolidation. Findings from both sleep-based and retrieval-mediated consolidation studies report comparable memory changes, including memory accuracy gains (Diekelmann et al., 2009; Hebscher et al., 2019), semanticization (Lifanov et al., 2021; Schapiro et al., 2017), long-term stability (Guran et al., 2020; Lutz et al., 2017), and shifts from hippocampal to neocortical areas (Ferreira et al., 2019; McClelland et al., 1995). Several studies have investigated sleep and retrieval training interventions in combination. These studies typically involve retrieval training followed by sleep, and test memory consolidation across both interventions using the same stimuli (Antony & Paller, 2018; Ashton et al., 2022; Mak & Gaskell, 2023; Schreiner et al., 2018; Zhang et al., 2025). However, if both interventions act upon the same memories, this approach for comparison introduces ceiling effects, making it difficult to separate their impacts on memories. To better compare their impacts on memory, it would be useful for sleep and retrieval training to act upon separate stimuli, in a counterbalanced order, and separate the interventions sufficiently in time to minimise crossover effects. Denis and colleagues (2024) have followed this approach by studying retrieval training and sleep interventions in separate conditions, but did not include a restudy control group, making it difficult to isolate if memory changes occur due to retrieval training or memory practice in general. Protocols that do not confound retrieval training and sleep, and employ appropriate control groups, would be useful when comparing brain-based mechanisms of memory consolidation, but this has yet to be investigated.

A well-researched moderator of sleep-based consolidation is encoding strength. Various studies report that weakly encoded memories are typically prioritised for consolidation over strongly encoded memories (Baena et al., 2020; Bäuml et al., 2014; Denis et al., 2020; Drosopoulos et al., 2007; Dumay, 2016; Petzka, Charest, et al., 2021; Schapiro et al., 2017, 2018). This encoding-strength-dependent selective enhancement of memories is proposedly due to strongly encoded memories already being somewhat neocortically integrated, such that sleep-based consolidation cannot act upon them further (Himmer et al., 2017). Sleep-based memory reactivation and the transfer to neocortical stores is thought to be mirrored during retrieval-mediated memory consolidation (Antony et al., 2017). It is plausible that retrieval-mediated memory consolidation also selectively enhances weakly encoded memories. Studies have found that memories weakened by interference and directed forgetting are selectively enhanced or saved by retrieval training (Abel & Bäuml, 2016; Halamish & Bjork, 2011; Pastötter et al., 2011). Additionally, some research finds that pre-sleep testing to strengthen memories improved post-sleep accuracy (Denis et al., 2023; Plihal & Born, 1997; Schoch et al., 2017). However, these studies did not directly examine encoding strength. A comparison of encoding-strength-dependent selective enhancement across both sleep and retrieval training would inform our understanding of mechanisms underlying memory consolidation.

The enhancement of weakly encoded memories is also linked to neural markers of sleep-based consolidation. Sleep spindles are approximately 12–16 Hz oscillations of the EEG signal during NREM sleep, that are generated in temporal sequence with slow-wave oscillations and memory reactivations to facilitate the transfer of information into the neocortex (Rasch & Born, 2013; Staresina, 2024). Sleep spindle parameters (i.e., spindle frequency, density, amplitude) have been shown to moderate the enhancement of weakly encoded memories across sleep, with greater sleep spindle activity relating to a greater enhancement of weakly encoded memories over strongly encoded ones (Chatburn et al., 2021; Creery et al., 2015; Denis et al., 2021; Kumral et al., 2023; Petzka, Chatburn, et al., 2021). Studies investigating the interaction between sleep spindles and encoding strength have measured memory outcomes using recall accuracy due to its consistent detection of sleep-based consolidation impacts on memories (Diekelmann et al., 2009). However, as retrieval training uses memory recall to consolidate memories, when comparing to sleep an alternate measure is needed, such as recognition accuracy. This will avoid additional study-test matching benefits that retrieval training would have above sleep, which may have impacted previous work comparing sleep and retrieval training (Bäuml et al., 2014; Mak & Gaskell, 2023; Zhang et al., 2025). To support this, previous research has used recognition accuracy to index the selective enhancement of memories across consolidation (Baena et al., 2020, 2021; Chatburn et al., 2021).

To compare encoding-strength-dependent memory enhancements across sleep and retrieval training, an effective proxy must be explored, as sleep spindles are absent during wake. Sigma power has been used as an alternative for spindle parameters to investigate memory changes across sleep-based consolidation (Piosczyk et al., 2013), due to it capturing activity within the sigma band (i.e., ∼12–16 Hz), in which sleep spindle activity occurs. Sigma power has also moderated encoding-strength-dependent forgetting reflecting contextual reinstatement during sleep (Schechtman et al., 2023), which also drives memory changes across retrieval-mediated consolidation (Antony et al., 2017). Some have speculated on the role of sigma-band activity during wake (e.g., Dringenberg, 2019), with Schreiner & Staudigl (2020) even suggesting it could be related to suppression of distractions as spindle-related neurons in the thalamus were associated with such during wake (Z. Chen et al., 2016). However, no study to date has explored a potential role for sigma power during wakeful memory consolidation.

Overall, if retrieval training and sleep induce the same brain-based mechanisms, it is expected that both consolidation interventions will only enhance weakly encoded memories when sigma power is high. The current study will test subjects across four conditions separated by at least 48 hr (retrieval training, restudy, sleep (120-min afternoon nap), and wake) and, using object-word pairs, assess immediate (encoding strength) and delayed (memory outcomes) recognition accuracy. Apart from the wake condition, EEG will used to estimate sigma power during the intervention period.

## 2. Methods

### 2.1. Subjects

Thirty individuals (27 women, 1 non-binary, 2 men) aged 18–34 (*M*=22.17, *SD*=4.21) participated in the study. Twenty-five completed all four conditions (retrieval, restudy, sleep, and wake), one subject only completed retrieval, one subject completed restudy, one subject completed wake, and two subjects completed sleep and wake conditions. One subject’s sleep condition was removed for poor EEG signal quality during the nap. EEG recording errors during the intervention period led to two restudy and three retrieval training conditions also being excluded (final sample: retrieval *n*=23, restudy *n*=24, sleep *n*=26, and wake *n*=28). The sample size was based on similar previous studies (Baena et al., 2020; Chatburn et al., 2021; Creery et al., 2015; Denis et al., 2021; Johnson et al., 2019; Oudiette et al., 2013; Petzka, Charest, et al., 2021; Schreiner et al., 2018; Zeng et al., 2021a). Owing to the nature of the paradigm, subjects were fluent English speakers, had normal or corrected vision, and self-reported no hearing or sleep problems. Subjects also reported on interview and questionnaire right handedness, no diagnosed psychiatric or sleep impairments, and were not taking any medications or recreational drugs in the last 6 months impacting EEG (Massand & Bowler, 2013; Nicholls et al., 2013; Weymar et al., 2010). Subjects were recruited via flyers posted around the university campuses, social media, and an online subject recruitment system. Subjects who completed all four conditions received a $200 honorarium. The University of South Australia Human Research Ethics Committee granted ethics approval for this project (protocol no. 205130).

### 2.2. Materials

#### 2.2.1. Pittsburgh Sleep Quality Index (PSQI)

Sleep habits across the last month were screened using the PSQI (Buysse et al., 1989). The PSQI contains 19 items which are used to generate a total score where scores >5 indicate significant sleep difficulties, which was used as a cutoff for eligibility. The PSQI has demonstrated adequate internal consistency and high test-retest reliability (Backhaus et al., 2002; Buysse et al., 1989; Carpenter & Andrykowski, 1998).

#### 2.2.2. Morningness-Eveningness Questionnaire (MEQ)

The MEQ was used to screen out subjects with extreme circadian types (Horne & Östberg, 1976). The MEQ is a self-report questionnaire involving 19 multiple choice questions on sleep/wake habits. Subjects with scores ≤30 or ≥70 were excluded. The MEQ’s items have previously demonstrated a high internal consistency and high test-retest reliability (Li et al., 2011; Pišljar et al., 2019).

#### 2.2.3. Flinders Handedness Survey (FLANDERS)

Handedness was screened for using the FLANDERS (Nicholls et al., 2013). Subjects were instructed to select their hand preference for each of the 10 tasks (e.g., drawing). Selecting “right” corresponds to +1 to their overall score, “left” is –1, and “mixed” is 0. Overall scores between –10 to –5 indicate left-handedness, –4 to +4 indicate mixed-handedness, and +5 to +10 indicate right-handedness. Subjects with scores <5 were excluded. The Flanders has displayed high internal consistency between its items (Nicholls et al., 2013).

#### 2.2.4. Karolinska Sleepiness Scale (KSS)

The KSS was used to control for sleepiness in the analyses (Åkerstedt & Gillberg, 1990). The KSS is a single-question self-report measure where subjects rate their current sleepiness on a scale from 1–10. Answers range from “1 – extremely alert” to “10 – extremely sleepy, cannot keep awake.” The KSS has demonstrated higher predictive validity for various measures of alertness than any other sleepiness measure, including reaction time (*r*=.57), number of lapses (*r*=.56), and power spectral density in both alpha (*r*=.40) and theta (*r*=.38) bands (Kaida et al., 2006).

#### 2.2.5. EEG

EEG was measured across each experimental phase (except the wake condition and break), using a 64-channel passive electrode BrainCap with Ag/AgCl electrodes positioned according to the modified 10–20 system (Jasper, 1958). Electrooculographic (EOG) activity was measured via two electrodes embedded in the cap, positioned approximately 1 cm diagonally away from the outer canthus of each eye. Electromyographic activity was measured via three electrodes (two placed on the skin above each subglottic muscle, and one placed between the subglottic muscles), and electrocardiographic activity was measured by one electrode placed above the heart. Two BrainAmp amplifiers (DC system, sampling rate 500 Hz) were used in combination with BrainVision Recorder software for output recording. Impedances were kept below 10 kΩ throughout all experimental phases.

#### 2.2.6. Stimuli

##### 2.2.6.1. Object images

We randomly sampled 416 object images (104 per condition) from sets 2–5 of the Mnemonic Similarity Task (MST; Stark et al., 2019). Each object image has another object image in the same set that is classed as a similar lure (e.g., a bouquet of flowers in a vase, versus a bouquet of flowers tied in a ribbon). To make sure the similar object lures were discriminable from each other above chance level, we only included MST images where the similar-object lure was endorsed less than chance level (i.e., ≤33.3%) in the MST’s accompanying dataset.

##### 2.2.6.2. Word sounds

Word stimuli were taken from an updated version of the Affective Norms for English Words (Warriner et al., 2013). This dataset contains 13915 English words, optimised for use in paired associate tasks, rated on frequency from 0–1000 and 1–9 on valence, arousal, and dominance. This study randomly selected 416 monosyllabic nouns (104 per condition), with valence and arousal ratings between 3.5–6.5. These words were recorded, using the voice of a native English speaker with an Australian accent, for their auditory presentation. These recordings were approximately 500–1000 ms long, each. Words were presented at approximately 70 dB.

### 2.3. Procedure

74 individuals expressed interest in participation, and were screened for their eligibility using the PSQI, MEQ, and FLANDERS (ineligible=12, loss of contact=32). The 30 eligible subjects then participated in four conditions, each of 4.5 hr (12:00–16:30; see Fig. 1), separated by at least 48 hr. The night before each condition, subjects were asked to wake up 1 hr earlier than usual to increase the likelihood of an afternoon nap entering NREM sleep, in the sleep condition. Subjects were seated in front of a dedicated computer and were fitted with an appropriately sized EEG cap. Subjects were then given details about the memory task they performed across the rest of the session.

**Fig. 1.**
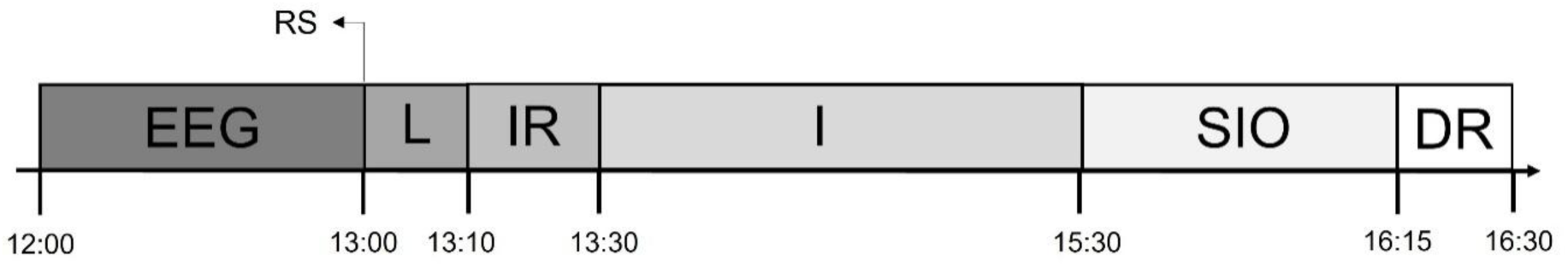
Procedure timeline. EEG = Setup EEG, RS = Resting State, L = Learning, IR = Immediate Recognition, I = Intervention (retrieval, restudy, sleep, or wake), SIO = Sleep Inertia Offset / Retention Period, DR = Delayed Recognition.

The memory task was based on previous protocols which have employed retrieval training and sleep (Ferreira et al., 2019; Liu et al., 2021; Schreiner et al., 2018). Open Sesame v.3.3.10 software was used to create and run each phase of the memory task (Mathôt et al., 2012). The memory task began with a learning phase, then immediate recognition, one of four intervention conditions (retrieval, restudy, sleep, or wake), and a final delayed recognition. The intervention conditions across sessions were experienced in a pre-determined, counterbalanced order. This counterbalancing was random, except that the subjects always completed one wakeful condition before the sleep condition, to increase their familiarity with the environment and therefore increase their likelihood of napping. The paired associates used in the memory task consisted of randomly paired MST object pictures and auditory words. All subjects were presented with the same object-word pairs, in a randomised order for each condition by subject. A different set of object-word pairs were used in each of the four conditions. For the object in each object-word pair, there was an object in a different object-word pair that was its similar object from the MST. Highly related object-word pairs were manually identified and reshuffled. A visualisation of the memory task can be found in Fig. 2. The following sections detail the memory task protocol.

**Fig. 2.**
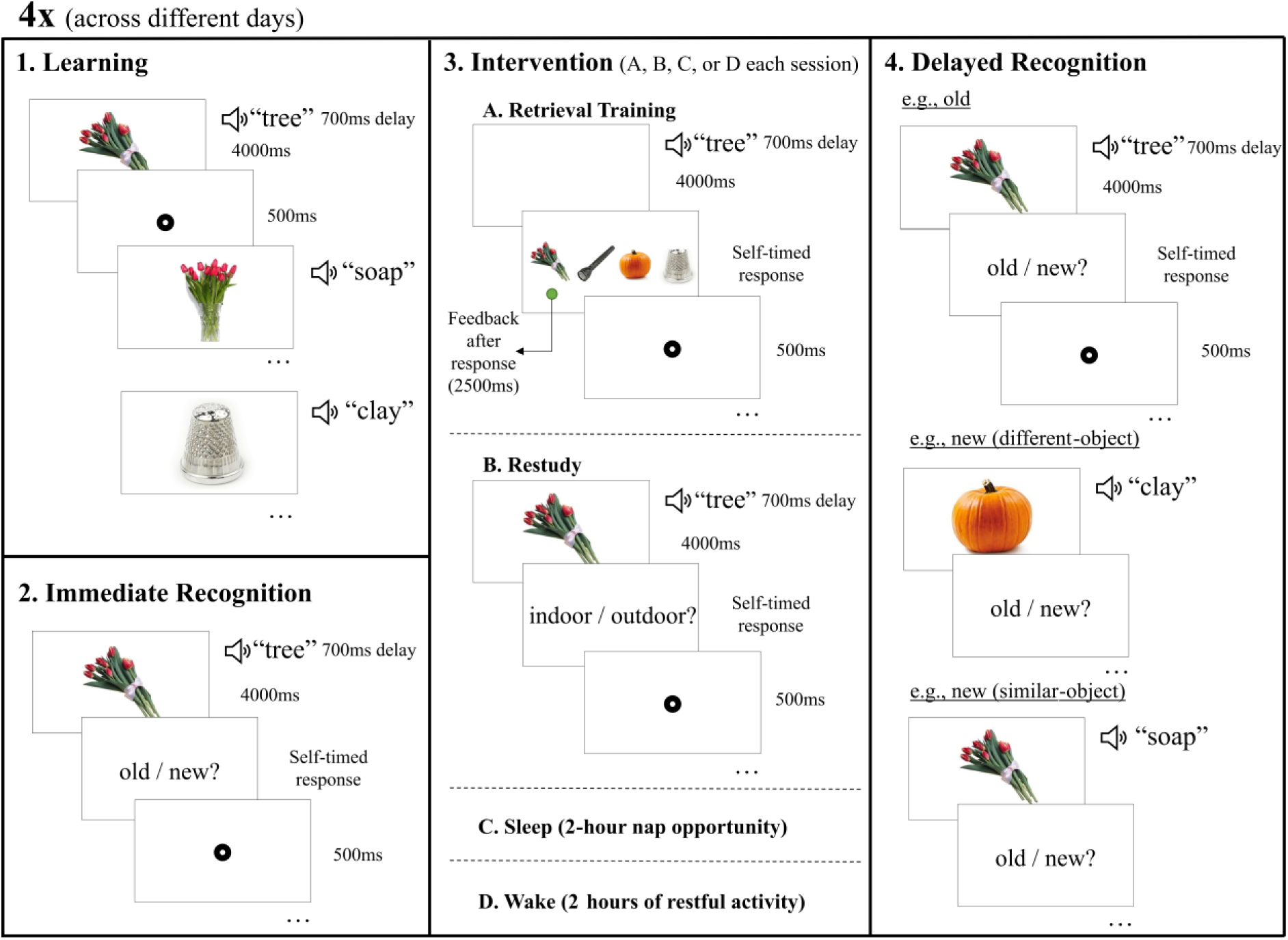
Memory task. 1. In the learning phase, subjects were exposed once to each of the object-word pairings. 2. Subjects then underwent immediate recognition testing, to distinguish between original (old) and rearranged (new) pairings, based on the pairings learnt during learning. An example of an old pairing is shown here. 3. Subjects underwent one of four intervention conditions. A. Involved retrieval training, where subjects heard each word, recalled the object, then picked it from distractors with feedback after. B. Subjects were exposed to all the pairs again and had to indicate if the object belonged indoors or outdoors. 1. C. Subjects had a 120-min nap opportunity. D. Subjects engaged in light activities. 4. Delayed recognition involved the same procedure as immediate recognition, where subjects distinguished between old and new pairings of the objects and words. This includes an example of an old pairing, new pairing with a different object to the object paired with the original word, and a new pairing with a similar object to the object paired with the original word.

#### 2.3.1. Learning

In each session, subjects learnt 104 object-word pairs. This involved viewing the image of an object on a white screen for 4000 ms and a sound of the paired auditory word was played 700 ms after the object onset. The images were displayed in colour, 400 pixels wide, in the centre of the screen. After each object was displayed, a fixation dot (8 pixels wide with a 2-pixel wide centre hole) was presented for 500 ms, before the next object was shown. Each pair was presented once.

#### 2.3.2. Immediate recognition

Immediately following learning, recognition memory of the pairs was tested by exposing subjects to each object and word again in the same sequence as the learning phase. This was done using an old/new paradigm, where 72 object-word pairs were presented in their correct pairings (old pairs), and 32 in a rearranged pairing (new pairs). To create the rearranged pairings, pairs were shuffled by substituting the object in each pair with another object of a different pair (16 similar objects, 16 different objects). After each pair was presented, a new white screen displayed the prompt “old or new?” Subjects responded via keypress, based on whether they believed that the pairing was presented as it was originally learnt (old / “Q”), or if it was rearranged (new / “P”). Before starting the task, subjects were encouraged to respond quickly, but optimize accuracy over speed. After they responded to the prompt, a fixation dot appeared on the screen for 500 ms before the next object-word pair was shown. No feedback was given to subjects on recognition accuracy performance. To avoid introducing interference effects with the original encoding episodes, the 32 rearranged (new) pairs were not used in subsequent phases.

#### 2.3.3. Intervention

For each of their four visits, subjects completed one of the four interventions, depending on the condition order they were assigned. Each intervention period took 120 min and EEG was recorded apart from one exception. This exception was the wake condition. After the intervention, all conditions included a 45 min retention period, which also served as a sleep inertia offset period for the nap condition. During this break, subjects remained in the laboratory and engaged in light activities (e.g., reading or scrolling on social media), simulating how they typically spend their free time. The KSS was completed at the end of the intervention. The four interventions are outlined below.

##### 2.3.3.1. Retrieval training

Subjects completed six rounds of retrieval training practice with the initially learnt object-word pairs. The 72 object-word pairs were randomly divided into two equal length lists, with each list being presented three times. Within each round, the order of presentation for the object-word pairs was randomised per subject. Practice round took 10−12 min. Subjects had 8–10 min breaks between each round so that the practicing was spaced out over the 120 min period.

The retrieval training practice had subjects hear the word of each pair, recall the paired object from their memory, then pick the object that was originally paired with the word from distractors. Subjects were presented a blank white screen and heard the word of one of the object-word pairs that were previously learnt. Subjects were instructed to imagine the paired object on the screen, when they heard the word. After 4000 ms, four object images that were part of previously learnt pairs were displayed 120 pixels wide, equally spaced across the width of the screen. Of the four objects, one of them was the original object that the word was paired with, while the other three were objects from other pairs. The distractor options were pre-determined and consistent across all subjects, but they were different each of the three times that a word was practiced. For every single pair that was practiced, exactly one round included the original object’s similar MST object as one of the distractors. Subjects were instructed to choose the object that they believed they previously learnt as paired with the word they just heard. Each distractor object had a black letter displayed above it which indicated the corresponding response key (Q, E, I, or P) required to select the object below it. 500 ms after their response, subjects received feedback on their selection via a green dot (8 pixels wide with a 2-pixel wide centre hole) placed below the correct object for 2000 ms. Subjects were instructed to use this feedback to learn the pairings and improve their memory. After the feedback was presented, a fixation dot was shown in the middle of the screen for 500 ms, before the next word was played on a blank screen. For the breaks between each round, subjects engaged in light activities.

##### 2.3.3.2. Restudy

In the restudy condition, subjects followed the same six-round study-break structure as the retrieval training intervention, practicing half (36) the 72 pairs at a time, three times each. Overall, restudy practice involved being exposed to both constituents of each word pair again. For each object-word pair, subjects were presented the object and word exactly as was done during learning, in a subject-specific randomised order. An object image from each pair was presented in the centre of the screen for 4000 ms, followed by the originally paired word presented auditorily 700 ms later. To maintain subjects’ attention, after each pair was presented, a screen prompted subjects to decide if the object of that pair would be commonly found indoors or outdoors. This prompt was displayed in the centre of a white screen, with the words “Indoors / Outdoors?” with the letter “Q” and underneath the word “Indoors,” and the letter “P” underneath “Outdoors.” These letters underneath the question depicted the keypress required to select the corresponding option. Subjects were instructed to treat this interactivity task as secondary to their practicing of the pairs. After subjects responded, a fixation dot appeared for 500 ms before the next object-word pair was presented.

##### 2.3.3.3. Sleep

Subjects were given a 120 min nap opportunity. Subjects slept in a quiet, darkened bedroom, and to promote sleep white noise was played at approximately 37db during the entire nap opportunity. The researcher remained in the adjacent room, monitoring the EEG signal for signs of sleep and any disturbances to the electrode signals.

##### 2.3.3.4. Wake

In the wake condition, subjects were supervised while they engaged in light activities (e.g., reading a book or watching a movie) for 120 min, to replicate their typical use of free time. During this time, subjects’ EEG was not recorded, but they remained in the laboratory with the EEG cap on.

##### 2.3.4. Delayed recognition

Comparable to the immediate recognition phase, subjects were presented with the same old/new task, but with a new assignment of pairs as either old or new. Subjects were exposed to the 72 object-word pairs they practiced, which were presented using the same sequencing as the immediate recognition phase. For half (36) of the 72 pairs, the word of each pair was presented with its originally paired object from the learning phase. The other half (36) of the object-word pairs were novel (new) pairings. Half of the novel pairings were similar-object lures, whereas the other half were different-object lures. In the exact same structure as the immediate recognition phase, subjects were prompted, by the screen following the object, to indicate if the pairing was old or new, receiving no feedback on their accuracy. After they answered, a fixation dot appeared for 500 ms before the next pair was presented. Whilst the order of presentation of pairs was randomised for each subject, all shuffled pairings and assignments to old or new categories were pre-determined to remain consistent across all subjects. After the delayed recognition test, subjects had the EEG cap removed, and then left the laboratory.

### 2.4. Data analysis

#### 2.4.1. Recognition accuracy (*d* prime)

Recognition accuracy in the immediate and delayed recognition tests was represented as *d* prime (*d*′), a sensitivity index of memory performance from Signal Detection Theory that accounts for false alarm rates when calculating accuracy (McNicol, 2005). The *d′* scores were calculated in *R*, v4.3.0 (R Core Team, 2022) using the *Psycho* (Makowski, 2018) package, by taking the difference between z transformed probabilities of hits (HI) and false alarms (FA) per subject (i.e., *d′*=z[HI] – z[FA]). Higher *d*′ scores represent a greater recognition memory of old pairs, controlling for potential high false alarm rates. Further, immediate recognition *d*′ scores represent a subjects average encoding strength across the pairs, whereas delayed recognition *d*′ scores represent their post-consolidation memory outcomes.

#### 2.4.2. EEG analyses

##### 2.4.2.1. EEG preprocessing

The EEG signals for the EEG intervention conditions (i.e., retrieval training, restudy, and sleep) were processed using *MNE-Python* v.1.7.0 (Gramfort et al., 2013). Activity from each intervention period was re-referenced to the mastoid electrodes M1 and M2 and down-sampled to 100 Hz (required for spindle detection). A notch filter (50 Hz) removed interference from the Australian mains power supply. Also, a finite impulse response filter using a Hamming window was applied to remove slow signal drifts below 1 Hz and high frequency activity above 40 Hz. Then, an Independent Component Analysis was applied to the retrieval training and restudy intervention’s data to correct electrooculographic (EOG) artifacts, excluding components which correlated the most strongly with EOG events (via the *create_eog_epochs* function in *MNE-Python*). Equal length 30-sec epochs were calculated across the six study periods for the retrieval training and restudy interventions, and during NREM sleep in the sleep intervention. EEG was segmented into 30 sec epochs for consistency across conditions, as this is the length of epochs used during sleep scoring. Sleep scoring of EEG data during the sleep intervention was performed as per established procedures (Berry et al., 2017).

##### 2.4.2.2. Individual Alpha Frequency (IAF)

Subjects’ IAF’s were used to create subject-specific frequency band limits for calculating time frequency representations. The IAFs were estimated using EEG data obtained from a 2-min eyes-closed resting state recording, during which alpha activity is typically most prominent (Grandy et al., 2013). This was represented by the centre of gravity, a weighted average of the power within the alpha band (∼7–13 Hz) that circumvents problems with identifying a dominant alpha peak. The alpha centre of gravity was calculated in MNE Python, using methods from Corcoran et al. (2018). Subjects’ IAF values were used to create subject-specific frequency bands for statistical modelling. This was done by normalising the upper and lower limits for frequency band bounds using the golden mean-based algorithm and their IAF, as described in Pletzer et al. (2010).

##### 2.4.2.3. Sigma power

Time frequency representations were calculated using a Morlet wavelet transformation (Cohen, 2019) within subject-specific sigma bands (∼12–16 Hz), for 5 cycles per 1 Hz frequency step. For each subject, the lower bound for their sigma band was defined as the upper bound for their alpha band, and the upper bound for each subjects’ sigma band was defined as the lower bound for their beta band. Absolute power was calculated, across each 30 sec epoch (NREM for sleep, during study for retrieval training and restudy conditions), per electrode and sampled time point. These power values were averaged across electrodes, time, and epochs, so that each subject session (one visit to the laboratory per subject) had a single value of average sigma power experienced across the intervention, measured in μV^2^/Hz. To determine regions of interest for the subsequent modelling, we averaged sigma power across topographical regions. These regions divided the electrodes into sagittal planes (i.e., anterior, central, posterior) from front-to-back, and lateral planes (i.e., left, midline, right) from left-to-right. This created a grid of nine intersecting topographical regions.

##### 2.4.2.4. Sleep spindles

The offline detection of sleep spindles in the sleep intervention condition, and the calculation of their parameters (i.e., density, frequency, and amplitude), was done via established methods in *MNE-Python* (Klinzing et al., 2016) using the *YASA* toolbox (Vallat & Walker, 2021). The data was first narrowed to NREM epochs and a subset of centro-parietal electrodes (Cz, C3, C4, Pz, P3, P4) commonly associated with fast spindles (Zeitlhofer et al., 1997). Individual peak spindle frequencies were identified per subject, to customise the detection of spindles events within 2 Hz either side of each subject’s peak. To detect spindles, a sliding window of 200 ms was used as a threshold to compute root-mean-square values. If the root-mean-square exceeded 1.5 standard deviations of the filtered signal for 500–3000 ms, the window was considered a sleep spindle (Lacourse et al., 2019). From these spindles, the density (number of spindle events divided the epoch length), frequency (median hertz of the spindles), and amplitude (peak-to-peak microvolt amplitude of detrended spindles) were calculated per epoch, channel, and sleep stage. These values were then averaged to obtain one value of each metric per subject’s condition.

#### 2.4.3. Statistical analyses

##### 2.4.3.1. Full sample

To test various predictions and interactions between variables, linear mixed-effects models were conducted using *R* v.4.3.0 (R Core Team, 2022) using the *lme4* package (Bates et al., 2015). Outliers were removed before each model was conducted; values were determined to be outliers if they were more than 1.5 times the interquartile range above quartile 3, or below quartile 1. As sigma power values and recognition accuracy are on different scales, all fixed effects and outcomes in the models were scaled for their effective comparison. This was achieved by subtracting the mean from all values (using the *scale* function from base *R*) and was applied to every model for consistency of interpretation. To determine if significant differences existed between the conditions within each model, we used treatment contrast coding. To plot the significant main and interaction effects from the linear mixed-effects models, the modelled effects were extracted using the *effects* package (Fox et al., 2022). In these plots, the error bars depicted the 83% confidence interval, which reflects a 5% significance threshold for non-overlapping estimates (Austin & Hux, 2002; MacGregor-Fors & Payton, 2013).

The first linear mixed-effects model aimed to predict delayed recognition accuracy from all four intervention conditions, controlling for immediate recognition accuracy. This was performed to gage the general impacts of all four conditions on recognition accuracy, as the wake condition could not be included in the subsequent modelling. This model used *d*′ scores at delayed recognition as the outcome, predicted by the fixed effect of intervention condition (i.e., retrieval training, restudy, sleep, and wake). This model controlled for *d*′ scores at immediate recognition to control for potential fluctuations (see Sassenhagen & Alday, 2016) and KSS scores at delayed recognition to control for the effects of sleepiness on one’s behavioural performance. Lastly, subject ID was used as a random effect.

The second model was conducted to determine the topographical region/s of interest for sigma power, to be used in the subsequent modelling to test our hypothesis. We took this data-driven approach due to the minimal research conducted on sigma power during wakeful memory tasks to inform a theory-driven region of interest. Only the three EEG intervention conditions (i.e., retrieval training, restudy, and sleep) were included in this model and the subsequent models, as no EEG was recorded during the wake intervention. This model predicted sigma power, using the fixed effects of EEG intervention condition, sagittal planes, and lateral planes. Subject ID was also used as a random effect. Treatment contrast coding was also applied in this model to determine significant differences within the model between the sagittal and lateral planes. As the role of sigma power during consolidation is not fully understood, we used a two-tailed approach for identifying significant regions (*p*<.025).

Finally, linear mixed-effects models were conducted to test the primary prediction of the study, that greater sigma power would increase the enhancement of weakly encoded memories across the intervention period, across retrieval training and sleep only. These models used *d*′ scores at delayed recognition as the outcome, predicted by the fixed effects of the EEG intervention conditions, *d*′ scores at immediate recognition (encoding strength), and sigma power. Additionally, this model controlled for delayed recognition KSS scores, and subject ID was used as a random effect. Each model averaged sigma power across one of the sigma power regions of interest identified in the prior modelling.

##### 2.4.3.2. Sleep subgroup

A series of exploratory models were conducted on only the nap condition’s data (*n*=26). The first set of models aimed to explore if sleep spindle parameters moderated the enhancement of weakly encoded memories. These models were performed to see if our study could replicate the findings of previous research, and therefore support any relationships observed with sigma power. Three linear models were conducted to predict *d*′ scores at delayed recognition as the outcome, using *d*′ scores at immediate recognition (encoding strength) as a predictor and the KSS scores at delayed recognition as a covariate. Each of the three models then included a different sleep spindle parameter as a predictor; density, frequency, or amplitude. These models were designed to mirror the first full sample model, with the only difference being the ID was not used as a random effect, as the nap group alone did not contain multiple values per subject.

Finally, exploratory models were conducted to give further evidence to whether sigma power was an effective proxy for sleep spindle parameters. To do this, Pearson correlations were conducted to explore if any of our sleep spindle parameters correlated with sigma power, during NREM sleep. Again, each correlation incorporated one of the three sleep spindle parameters; density, frequency, or amplitude. These correlations were run for all sigma power regions of interest that were identified in former modelling. Bonferroni corrections were applied to these correlations by dividing the significance threshold by the number of correlations run (e.g., *p*<.008 for six correlations across two regions of interest).

## 3. Results

The subject-specific sigma bands in this sample had a mean lower bound of 12.40 (*SD*=1.48) Hz and a mean upper bound of 16.30 (*SD*=1.93) Hz. Table 1 displays the sleep parameters of subjects during the nap intervention. All subjects experienced NREM sleep during their nap opportunity, and only five subjects did not enter slow-wave sleep. Four subjects demonstrated rapid eye movement (REM) sleep during their nap. No subject experienced sleep onset REM.

**Table 1.** Sleep characteristics of the nap intervention.

| Sleep Measure | Mean (SD) | Range |
| --- | --- | --- |
| TST (min) | 82.44 (26.31) | 25.00–116.00 |
| SOL (min) | 11.90 (11.11) | 1.50–53.00 |
| N1 (%) | 16.70 (9.54) | 5.00–49.50 |
| N2 (%) | 43.90 (14.00) | 19.50–71.00 |
| SWS (%) | 19.00 (18.31) | 0.00–56.00 |
| REM (%) | 2.87 (6.30) | 0.00–20.50 |

*Note.* TST = total sleep time, SOL = sleep onset latency, N1 = stage 1 non-rapid eye movement (NREM) sleep, N2 = stage 2 NREM sleep, SWS = slow-wave sleep, REM = rapid eye movement sleep.

### 3.1. Recognition accuracy improvements were greatest after retrieval and restudy interventions

Table 2 depicts the *d*′ recognition accuracy scores at immediate and delayed recognition. These values indicate that recognition accuracy improved somewhat from before to after each intervention.

**Table 2.** Recognition accuracy *d′* scores across each condition.

| Intervention Condition | Mean Immediate Recognition $d'$ (SD) | Mean Delayed Recognition $d'$ (SD) |
| --- | --- | --- |
| Retrieval | 0.93 (0.78) | 2.49 (0.90) |
| Restudy | 0.95 (0.81) | 2.20 (1.06) |
| Sleep | 1.01 (0.77) | 1.42 (0.91) |
| Wake | 0.75 (0.50) | 0.94 (0.59) |

*Note.* Mean values are reported across subjects, per condition. SD=standard deviation, which is reported in brackets.

Additionally, a linear mixed-effects model tested for delayed recognition accuracy differences between all four intervention conditions. As differences in immediate recognition accuracy were observed between conditions (Table 2), this was controlled for in the model. The model found a significant main effect of condition to predict delayed recognition accuracy, *χ^2^*(3)=129.59, *p*<.001. Based on Fig. 3, delayed recognition accuracy was greatest after retrieval training and restudy, compared to sleep and wake interventions. Treatment contrast coding revealed various significant differences that are depicted in Fig. 3.

**Fig. 3.**
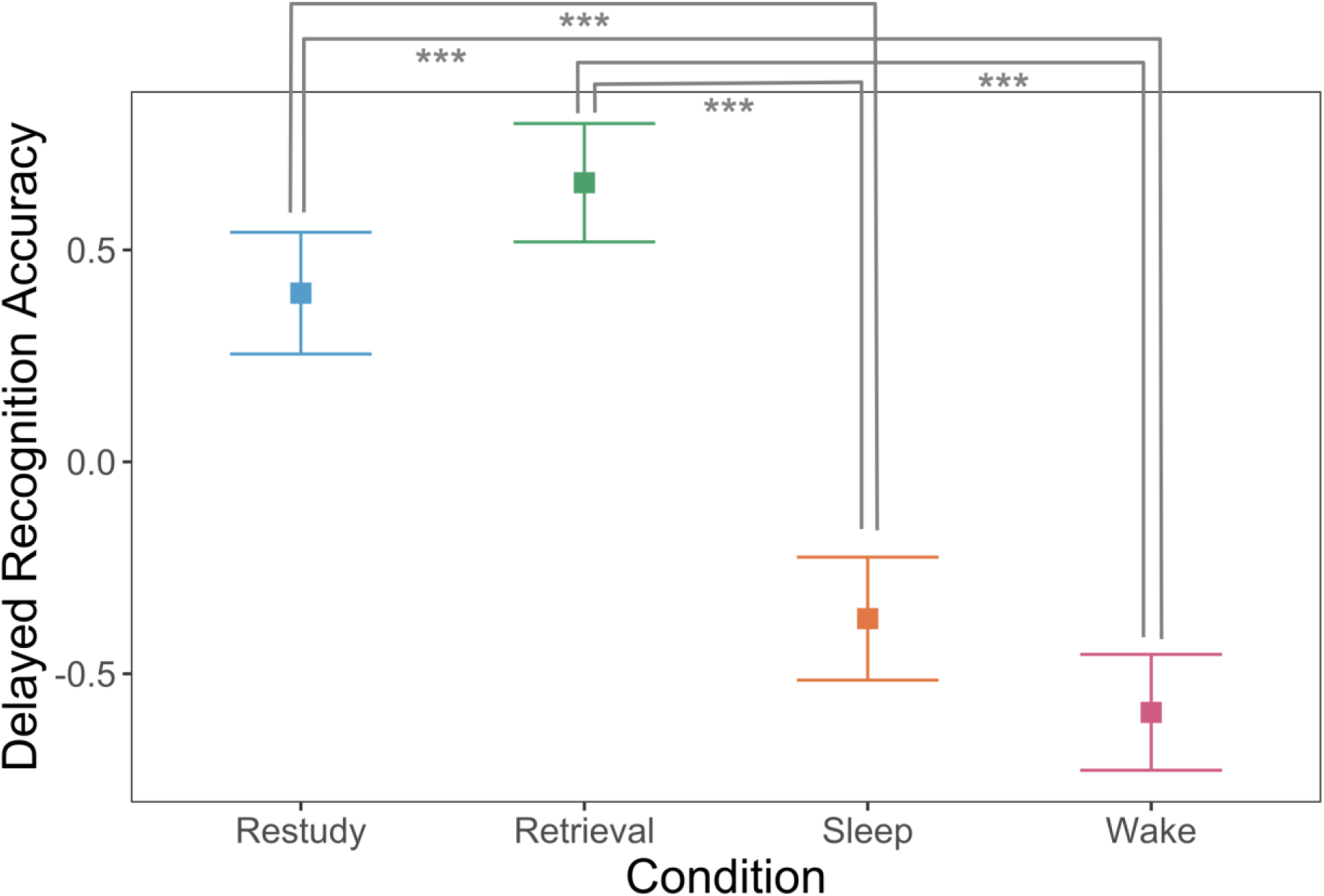
The x-axis represents modelled and centred delayed recognition accuracy, measured via *d* prime. The error bars represent the 83% confidence intervals. *denotes *p*<.05, \*\**p*<.01, and \*\*\**p*<.001.

### 3.2. Encoding strength and posterior sigma power predict delayed recognition accuracy

To determine topographical regions of interest for sigma power, we conducted a linear mixed-effects model predicting sigma power from the intervention condition, as well as the sagittal and lateral planes that the sigma power was averaged across. This model revealed a significant interaction between sagittal and lateral planes to predict sigma power (*χ^2^*(4)=19.18, *p*<.001), as well as an interaction between condition and sagittal planes (*χ^2^*(4)=22.02, *p*<.001). Fig. 4 depicts both interaction effects. Subsequent treatment contrast coding revealed that sigma power was greatest in anterior and central midlines regions, across all conditions. This testing also revealed that sigma power was lowest overall in posterior regions for the sleep condition. Therefore, the subsequent modelling will investigate two regions of interest across all conditions, an anterior/central midline region and a posterior region.

**Fig. 4.**
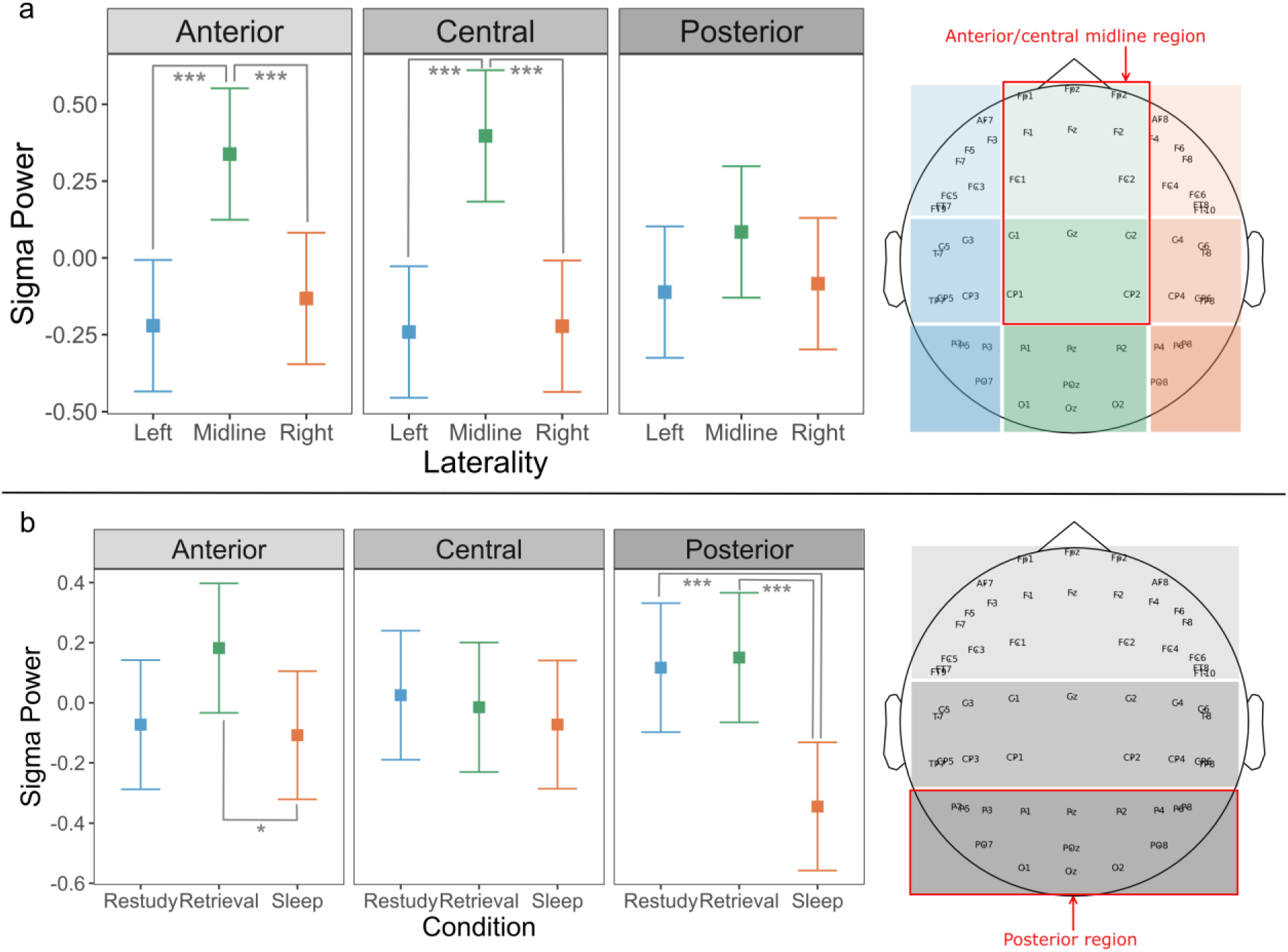
Differences in sigma power across conditions, sagittal planes, and lateral planes. Across all subplots, sigma power is centred around 0 and reflects the modelled effects from the linear mixed-effects model. The scalp plot on the righthand side of both subplots depicts the EEG channel locations and their respective sagittal and/or lateral cross regions. The error bars represent the 83% confidence intervals. Significant treatment contrasts are marked as: *denotes *p*<.05, \*\**p*<.01, and \*\*\**p*<.001. a) The interaction between sagittal and lateral planes to predict sigma power. The scalp plot identifies the anterior/central midline region of interest that is used in subsequent modelling. b) The interaction between sagittal planes and condition to predict sigma power. The red box on the scalp plot identifies the posterior region of interest for use in subsequent modelling.

A linear mixed-effects model was conducted to detect if anterior/central midline sigma power moderates an enhancement of weakly encoded memories over strongly encoded ones. This model found no evidence for a significant interaction between anterior/central midline sigma power, immediate recognition *d*′ (encoding strength), and EEG intervention condition to predict delayed recognition *d*′, *χ^2^*(2)=1.16, *p*=.561. The model did reveal two significant main effects of immediate recognition *d*′ (*χ^2^*(1)=61.04, *p*<.001), condition (*χ^2^*(2)=39.79, *p*<.001) to predict delayed recognition *d*′. The condition main effect is captured by Fig. 3, Fig. 5 depicts the main effect of immediate recognition *d*′ on delated recognition *d*′. No further main or interaction effects were observed between any predictors in the model.

**Fig. 5.**
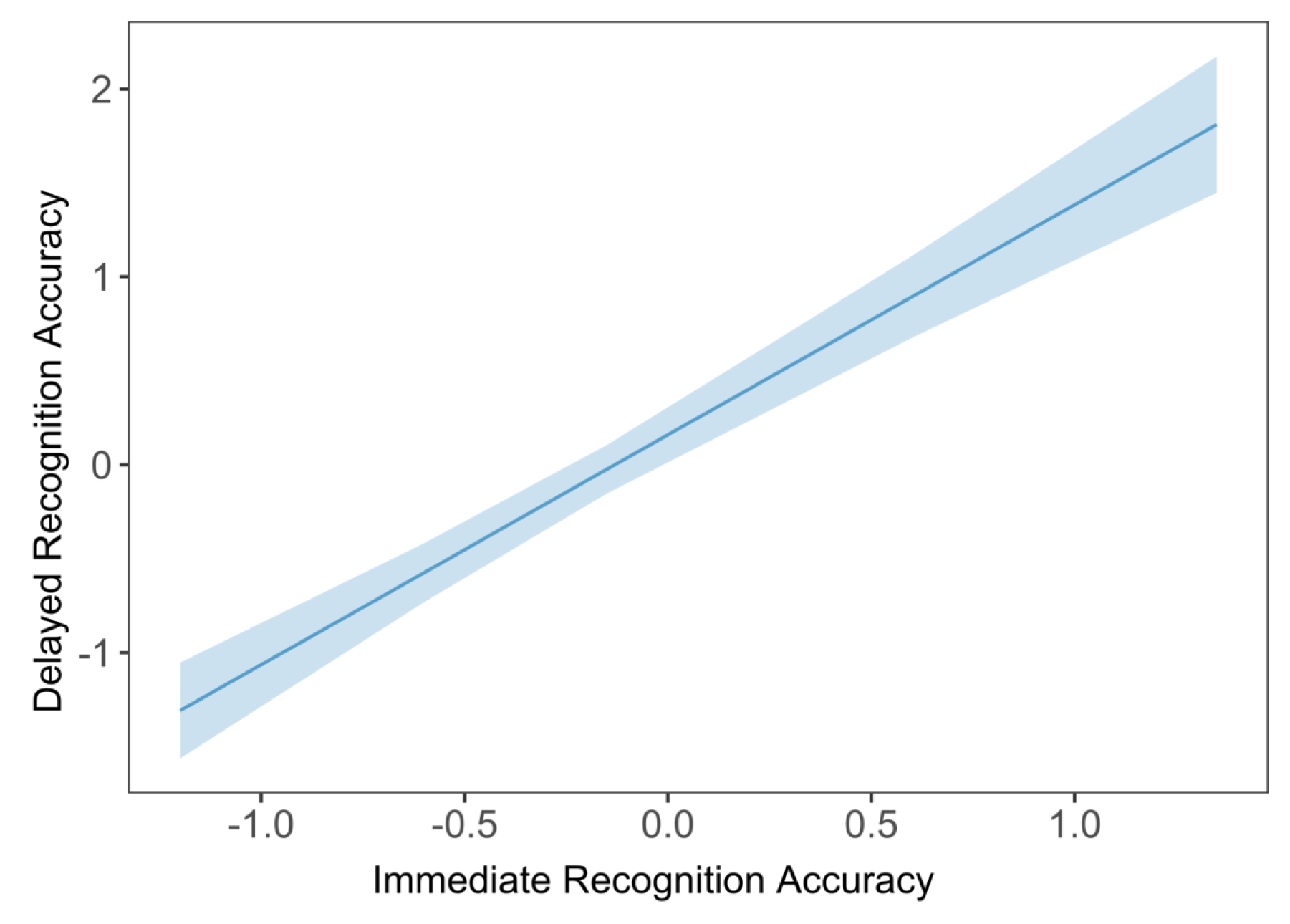
Main effect of encoding strength to predict delayed recognition accuracy. The main effect of immediate recognition accuracy (encoding strength) to predict delayed recognition accuracy. Immediate and delayed recognition accuracy are both measured as *d* prime, reflecting the modelled effects from the linear mixed-effects model.

A similar linear mixed-effects model as above was performed to identify if posterior sigma power moderates an enhancement of weakly encoded memories over strongly encoded ones. This model similarly found no evidence for a significant interaction effect between all predictors (*χ^2^*(2)=0.91, *p*=.633) and was able to replicate the significant main effects of condition (*χ^2^*(2)=48.78, *p*<.001) and encoding strength (*χ^2^*(1)=77.92, *p*<.001) to predict delayed recognition accuracy. In contrast to the model above, this model additionally detected a significant main effect of posterior sigma power to predict delayed recognition accuracy (*χ^2^*(1)=4.02, *p*=.045), which is illustrated in Fig. 6. No further main or interaction effects were observed between any predictors in the model. To determine if this effect was present in the posterior regions of the theta, alpha, and beta bands instead, exploratory modelling was conducted. We replicated the model above thrice, with each model replacing posterior sigma power with either posterior theta, alpha, or beta power. All three exploratory models detected no significant main or interaction effects involving EEG power.

**Fig. 6.**
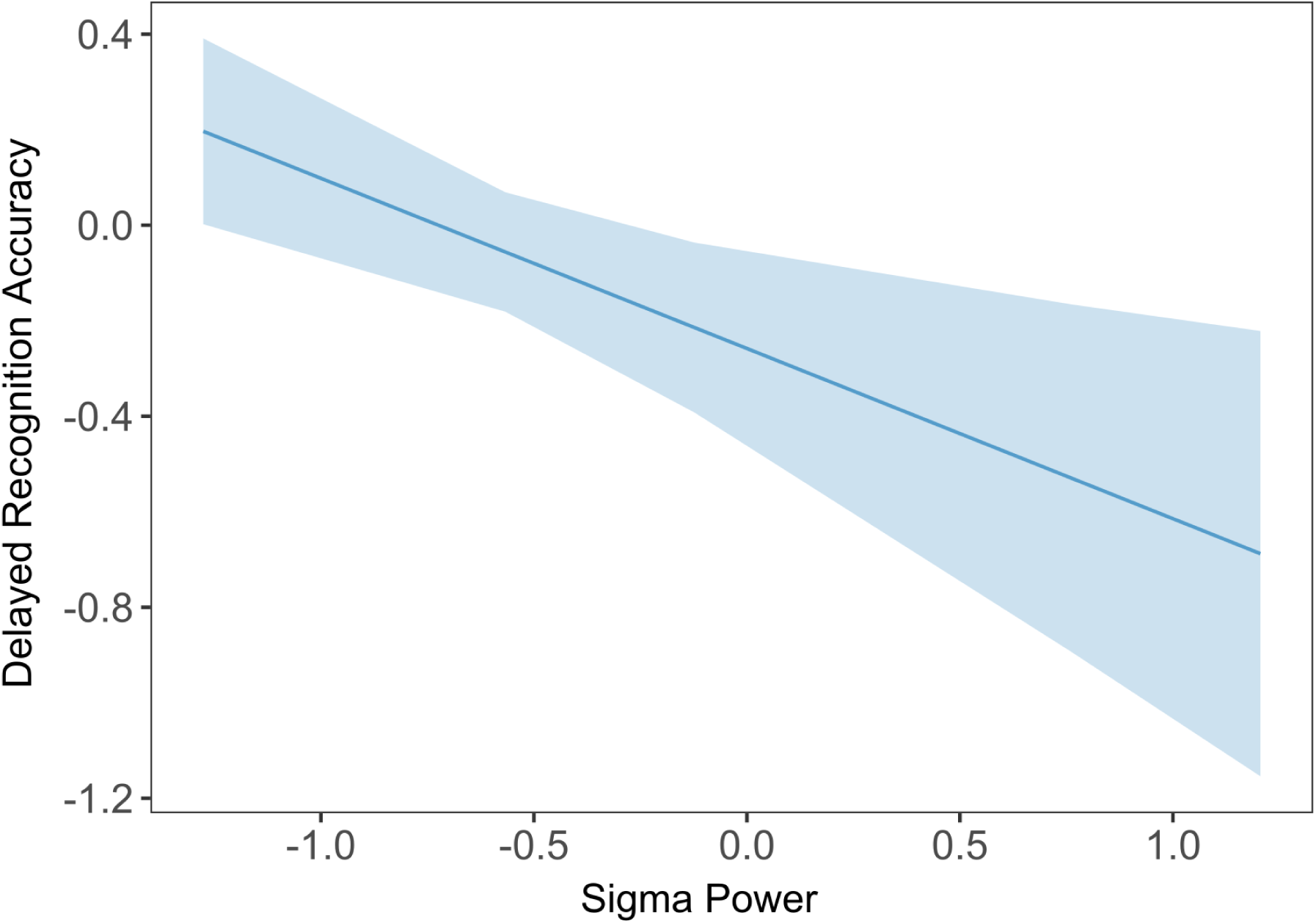
Main effect of posterior sigma power to predict delayed recognition accuracy. The main effect of sigma power to predict delayed recognition *d′*. Sigma power is centred around 0. Delayed recognition accuracy is measured as *d* prime, reflecting the modelled effects from the linear mixed-effects model.

### 3.3. Sleep spindle amplitude correlated with sigma power, but did not predict memory outcomes

Additional exploration was performed on the sleep group only, to reproduce previous findings that sleep spindle parameters moderate weakly encoded memories. This was tested using three linear models, replicating the sigma power linear mixed-effects models described above. Each of these models predicted delayed recognition *d*′ from immediate recognition *d*′ (encoding strength), with either spindle density, frequency, or amplitude instead of sigma power as the other predictor. However, no significant interaction effect was found between immediate recognition *d*′ and either spindle density (*F*(1)=0.93, *p*=.346), frequency (*F*(1)=0.25, *p*=.622), or amplitude (*F*(1)=0.53, *p*=.471), to predict delayed recognition *d*′. These models also did not reveal any main effects of spindle parameters to predict delayed recognition outcomes.

We then tested if both the anterior/central midline and posterior regions of sigma power were related to each spindle parameter, and if they therefore served as an effective proxy measurement. To accomplish this, we conducted three Pearson correlations with each sigma power and each of the three spindle parameters (i.e., spindle density, frequency, and amplitude), for both the anterior/central midline region and the posterior sigma power regions. The results of these correlations are reported in Table 3, which shows that only sleep spindle amplitude positively correlated with anterior/midline sigma power and posterior regions of sigma power.

**Table 3.** Results from Pearson correlations between sleep spindle parameters and sigma power regions.

| Sigma power region | Sleep spindle parameter | *df* | *r* | *p* |
| --- | --- | --- | --- | --- |
| Anterior/central<br>midline | Density | 24 | 0.29 | .140 |
|  | Frequency | 24 | 0.08 | .690 |
|  | **Amplitude** | **24** | **0.77** | **<.001** |
| Posterior | Density | 24 | 0.19 | .370 |
|  | Frequency | 24 | 0.42 | .039 |
|  | **Amplitude** | **24** | **0.54** | **.005** |

*Note.* *df* = degrees of freedom. Bonferroni corrected significance cutoff is $p < .008$ . Significant correlations are in bold.

## 4. Discussion

The broad goal of this study was to test if memory consolidation has a consistent brain-based mechanism to selectively enhance memories across sleep and retrieval training. To test this, we investigated if both sleep and retrieval training interventions enhanced weakly encoded memories over strongly encoded memories, and the role played by different topographical regions of sigma power. We also attempted to replicate the enhancement of weakly encoded memories across sleep, moderated by sleep spindle parameters. Contrary to previous research, we did not detect an enhancement of weakly encoded memories, moderated by either sigma power or sleep spindles. Intervention condition, encoding strength, and sigma power did not interact to predict memory outcomes. Nonetheless, we did find that intervention condition, encoding strength, and posterior, but not anterior/central midline, sigma power predicted recognition accuracy, independently. Additionally, we found that sigma power correlated with sleep spindle amplitude, but not frequency or density.

Consistent with the expectations that practice has the greatest impact on memory outcomes, retrieval training and restudy conditions produced significantly higher delayed recognition accuracy than the sleep and wake interventions. This is in line with research that retrieval training and restudy similarly boost initial memory accuracy, but the benefits of retrieval training over restudy typically only appear after longer delays (Bäuml & Trißl, 2022; Ferreira et al., 2019; Ferreira & Wimber, 2023; Guran et al., 2020). Whilst there was still an enhancement of recognition accuracy scores across the sleep and wake interventions, the magnitude of this was far smaller than either practice intervention. This is not entirely surprising, as greater exposure to memories, and the ability to update them, leads to a better recognition accuracy (Hintzman, 1970). Instead, the slight enhancement of memories across sleep is more in line with research that sleep typically protects memories from short-term decay (Denis et al., 2023; Dumay, 2016; Rasch & Born, 2013).

It is important to consider that condition differences in recognition accuracy do not speak entirely to a difference between sleep-based and retrieval-mediated consolidation mechanisms. This is because recognition accuracy alone does not give absolute insight into consolidation mechanisms or the qualitative changes it imposes on memories. Rather, it is typically the context of behavioural scores that reflect consolidation more reliably (e.g., considering encoding strength and sigma-band activity). More direct comparisons of sleep-based and retrieval-mediated consolidation with other behavioural markers and paradigms (e.g., lure endorsement, recall accuracy, or 1-week follow up testing) will be useful to see if any similarities or differences exist in how they uniquely reflect memory changes. These comparisons will more effectively speak to the fine-grained nature of memory changes across consolidation.

We were unable to detect an enhancement of weakly encoded memories across retrieval training or sleep conditions. Instead, we detected a positive relationship between encoding strength and subsequent memory outcomes, regardless of condition. Various studies also report that strongly encoded memories are prioritised for consolidation instead of weaker ones, and that this is also moderated by sleep spindle parameters (Schoch et al., 2017; Tucker & Fishbein, 2008; Wernette & Fenn, 2024; Wislowska et al., 2017). These opposing findings are typically attributed to different studies exploring different halves of an inverted-U-shaped curve, wherein too weakly or too strongly encoded memories are not consolidated, and moderately encoded memories are instead prioritised (Stickgold, 2009). In our study, we expected that learning relatively simple paired associates would place subjects across the right-most half of the inverted-U-shaped curve, wherein weaker encoded memories are prioritised. Alternatively, it is likely that these results are independent from the effects of consolidation, and therefore have no place on either side of the inverted-U-shaped curve. In our study, the enhancement of strongly encoded memories was not moderated by sigma power or any sleep spindle parameters, and the observed relationship did not differ in the restudy and wake conditions where consolidation was not being induced. Instead, it is plausible that our subject-level measure of broad immediate recognition may have been influenced by subjects’ trait-like learning ability, intelligence, and/or motivation which reflect separate processes to encoding strength (Creery et al., 2015; Denis et al., 2020). Other studies have attempted to avoid this influence by experimentally manipulating encoding strength (Drosopoulos et al., 2007; Schapiro et al., 2017), but research has still found that allowing for a more naturalistic measure of encoding strength, such as ours, still detects these relationships (Chatburn et al., 2021; Djonlagic et al., 2009). Overall, this relationship was likely not due to consolidation-related changes, but rather trait-like influences of learning ability that do not speak to a commonality unique to sleep and retrieval training interventions.

The posterior region of sigma power during the intervention period was negatively related to memory outcomes, contrary to the beneficial role of sleep spindles on memory outcomes (Antony et al., 2018; Gibson et al., 2022; Petzka, Chatburn, et al., 2021; van der Helm et al., 2011). Even the anterior/central midline region, where greater sigma power occurred, did not detect this predicted relationship. Despite the sleep condition experiencing the lowest posterior sigma power, there was no condition difference for the impact of posterior sigma power on memory outcomes. This may mean that within-condition sigma power fluctuations between subjects were likely driving this relationship, and that the range of these fluctuations was simply lower overall in the sleep condition. Moreover, the negative relationship between posterior sigma power and memory outcomes cannot be explained by a multicollinear relationship between sigma power and condition, as the sleep condition involved lower sigma power than sleep and a lower delayed recognition accuracy, not higher. Regardless, for posterior sigma power to relate to memory outcomes without interacting with the intervention condition variable, means that sigma power may not reflect a common mechanism between sleep-based and retrieval-mediated consolidation. Instead, it may reflect a memory mechanism across multiple interventions and states. Previous research has identified a negative subsequent memory effect (i.e., reduced power for subsequently remembered items) in the alpha/beta bands, which is especially prominent in posterior regions (Fellner et al., 2013; Klimesch et al., 1996; Sederberg et al., 2003, 2006; Waldhauser et al., 2012). As the sigma-band frequencies (∼12–16 Hz) exist within the alpha/beta range (∼8–30 Hz), one may assume that we have captured a fraction of a more widespread effect. However, when we ran the models predicting delayed recognition accuracy from encoding strength again with posterior alpha (∼8–12 Hz) and beta (∼16–25 Hz) power, no significant relationship was identified for either frequency band. This could suggest that sigma-band activity may underlie these previous findings, which was attributed across the wider alpha-beta frequency range by previous research not accounting for the subject-specific variations in frequency bounds. This identifies a novel strength of our approach, as we were able to isolate a subject-specific effect using custom frequency bands, that other research had not found using canonical band limits. Future research should identify if subject-specific frequency ranges specifically contribute to this negative relationship between EEG power and memory outcomes across a variety of paradigms. This may help identify if subject-specific frequency variations and/or task-specific manifestations underlie why our results differ from previous research. Overall, the precise roles that sigma power plays are unclear at present. To greater understand the role of sigma power during memory interventions, future research should continue to identify in which contexts sigma power predicts memory outcomes, and if it plays a different role during consolidation or across different mental states.

One key aspect that may explain why we did not detect an enhancement of weakly encoded memories moderated by sigma power, is that we estimated sigma power across large epochs during key periods of the interventions (e.g., Piosczyk et al., 2013), rather than around specific reactivation events (e.g., Schechtman et al., 2023). Antony and colleagues (2018) found that only the spindle events that were locked to memory reactivations, predicted memory outcomes. Therefore, measuring sigma power and sleep spindles outside of reactivation events, whilst unavoidable in our paradigm, may have incorporated a large amount of noise in our measure. This may have abstracted our sigma power measurement beyond the point that it was able to reflect sleep spindle processes. This is supported by the fact that we were unable to detect a relationship between spindle density/frequency and sigma power, which are more commonly related to memory outcomes than sleep spindle amplitude (P. Chen et al., 2024; Kumral et al., 2023). Whilst we did find a relationship between sigma power and sleep spindle amplitude, this is likely driven by the fact that the sleep spindle amplitudes in centro-parietal electrodes contributed to the sigma power calculations in the overlapping regions of interest. Despite this relationship, sleep spindle amplitude did not correlate with memory outcomes. Additionally, we explored if theta, alpha, or beta power during the same 30-sec epochs predicted memory outcomes. These models did not detect any significant relationships between EEG power and memory outcomes, despite research linking power in these frequency bands to memory outcomes when power is locked to specific events (Aktürk et al., 2022; Hanslmayr et al., 2009; Klimesch et al., 2006; Vivekananda et al., 2021). Therefore, our measure of EEG power may not have been sensitive enough to the time-locked events that were driving the memory changes we aimed to measure, potentially leading to the non-detection of this complex relationship.

Despite previous research, we did not find a role of spindle parameters in enhancing weakly encoded memories. These models were unlikely to be underpowered, as similar sample and stimuli sizes have been used previously to detect effects in similar studies (Baena et al., 2020; Chatburn et al., 2021; Creery et al., 2015; Denis et al., 2021; Johnson et al., 2019; Schreiner et al., 2018; Zeng et al., 2021b). These results are also not likely to be influenced by age-related differences in sleep EEG, as our sample did not contain subjects over 35 years of age (Landolt & Borbély, 2001). Additionally, whilst our study used a nap rich in NREM opposed to a full night of sleep, two meta-analyses have found no significant differences in the impact of spindle activity on memories between naps and full-nights of sleep (P. Chen et al., 2024; Kumral et al., 2023). Instead, our inability to detect a role of spindles in this study could be due to consolidation not acting upon weakly encoded memories in such a way that it is detectable through recognition accuracy. Perhaps, the enhancement of weakly encoded memories may be more accurately described as an enhancement of the *recall accuracy* of weakly encoded memories. Consolidation across sleep and retrieval training may more specifically strengthen connections between weakly encoded memories and the retrieval cues, thereby increasing their accessibility, which is less necessary for the memory search undergone during recognition memory (Gillund & Shiffrin, 1984). Such a distinction is imperative for future research to investigate further by directly comparing recognition and recall accuracy across consolidation.

Our study pioneers a sound methodological approach for comparing retrieval training and sleep interventions. Most previous studies that compare these consolidation interventions, do so with confounding sequential designs that produce ceiling effects, or draw cross-paradigm comparisons despite the notorious paradigm-specific manifestations of consolidation-related memory changes (Cordi & Rasch, 2021; Diekelmann et al., 2009). Additionally, our use of control conditions for both retrieval training and sleep interventions, aids in ruling out general practice effects on memory. To better compare sleep and retrieval training interventions, it would be useful for future studies to take our methodological approach of examining these interventions in parallel and in the same paradigm. Our approach is one of the first to do this, and continuing these direct comparisons will inform a more comprehensive theory of memory consolidation that considers memory changes across multiple states and contexts. More specifically, to compare more fine-grained time-resolved EEG mechanisms between sleep and retrieval training, future comparisons should isolate EEG around key consolidation-related events and track the item-level changes of memories. Additionally, investigating if encoding strength impacts certain aspects of memory outcomes (e.g., recognition, recall, false alarms) more than others, will further identify the fine-grained memory qualities that retrieval training and sleep types prioritise during consolidation.

In summary, while our results did not demonstrate a role of sigma-band activity in enhancing weakly encoded memories across both sleep-based and retrieval-mediated memory consolidation, we did find a role for post-learning posterior sigma power to impact subsequent memory outcomes across both sleep and wakeful states. We also demonstrated that the enhancement of weakly encoded memories may not be ubiquitous enough to impact recognition memory. Most crucially, our study offers a successful method for comparing consolidation across different states, using parallel sleep and retrieval training interventions, and adequate control conditions for both, all across different sessions. Continuing to make similar comparisons in future will help understand these two consolidation types, as their impacts manifest in different paradigms.

## Acknowledgements

We would like to thank the subjects for their time.

## CRediT authorship contribution statement

Hayley B. Caldwell: Conceptualization, Data curation, Formal analysis, Investigation, Methodology, Project administration, Validation, Visualization, Writing – original draft, Writing – review and editing.

Kurt Lushington: Conceptualization, Methodology, Project administration, Resources, Supervision, Writing – review and editing.

Alex Charburn: Conceptualization, Methodology, Project administration, Resources, Supervision, Writing – review and editing.

